# Immunophenotypic changes in peripheral blood of colorectal cancer patients: a potential diagnostic marker for CD4+ monocytes

**DOI:** 10.1101/2024.08.26.609793

**Authors:** Khin Aye Thin, Andrew Cross, Phonthep Angsuwatcharakon, Apiwat Mutirangura, Charoenchai Puttipanyalears, Thomas Pike, Robert Jones, Paul Skaife, Steven W Edwards

## Abstract

Colorectal cancer (CRC) can induce stresses on the immune system that can affect both the numbers and function of these cells and the ability of the tumours to evade immune-surveillance. Changes in immune functions can also occur during ageing and these may affect both the ability to fight infections and to protect against cancers. As the incidence of CRC is age-related, the aim of this work was to identify changes in immune cell subtypes that are specific to CRC and not merely due to age-related changes. The immunophenotypes of peripheral blood lymphocytes and monocytes of CRC patients and age-matched healthy controls (HC) were analysed using flow cytometry and surface marker staining. Compared to HC, a lower L:M ratio was observed starting from the early stages of CRC, while numbers of B lymphocytes were lower and CD4+ monocytes were higher in CRC patients. In most patients with CRC, the numbers of helper T cells were lower, while cytotoxic T cells and NK T cells together with CD4+ NKT and CD8+ NKT cells were higher: classical monocytes were lower while intermediate monocytes were higher throughout the stages of CRC, and HLA-DR^low^ monocytes were also higher. NK^bright^ cells were higher while NK^dim^ cells were lower in patients with large tumours. Most of the increased numbers of T cells and monocytes in CRC relate to immunosuppressive phenotypes that may aid tumour evasion. The increase in CD4+ monocytes is likely related to increased numbers of intermediate monocytes, and a threshold of 11.6% CD4+ monocytes can be used as diagnostic marker for CRC with a 60% sensitivity and 88% specificity.

## Introduction

CRC is the third most common cancer and second highest leading cause of cancer deaths worldwide [1]. The incidence of the disease can vary with sex, age and geographical regions and is affected by lifestyle factors: incidence of disease is higher in males than in females and higher in more-developed regions than in less-developed regions [2]. Some CRC cases are hereditary, but the majority of CRC patients are of the sporadic type with no family history of the disease [3]. The major risk factor for CRC is ageing with the majority of sporadic CRC patients >50 years old at the time of diagnosis [4]: 75% of rectal cancer patients and 80% of colon cancer patients are in their 60s and above [3].

During ageing, there is increased genome-wide hypomethylation and accumulation of temporary DNA modifications such as epigenetic marks, DNA damage or lesions [5]. CRC is a heterogeneous disease also associated with chromosomal instabilities such as genetic mutations, epigenetic alterations and environmental factors which are all involved in pathogenesis [6]. Moreover, colonic epithelial proliferation increases during ageing [7] and cumulative molecular and pathophysiological changes in the homeostasis of colonic epithelial cells throughout life can lead to neoplasia in the elderly [6]. It is therefore important to distinguish molecular changes that are associated with the ageing process and those that are associated with development of cancer.

Immune cells are dynamic populations of cells [8] and changes in their numbers, function and activity can be affected by: (a) unmodifiable factors such as age, gender; (b) modifiable factors such as stress, exercise and diet and (c) pathological factors such as inflammation and diseases [9–13]. Colorectal cancer can result in chronic stresses to the immune system and can also contribute to changes in the functions of immune cells [14]. However, circulating immunophenotypes change during ageing [15]. For example, helper T cells increase while cytotoxic T cells decrease [16]; NK cells expand in old age [17]; classical monocytes decrease while intermediate monocytes increase [18].

Therefore, the aim of this study was to identify immunophenotypic changes of subsets of T cells, NK cells and monocytes in CRC that may indicate altered functions as a result of cancer and may be useful as diagnostic markers of disease. It was important to characterize any changes in these phenotypes that were a function of disease (CRC) and not merely a consequence of ageing. We show increased numbers of suppressive immune phenotypes in CRC that may aid tumour evasion.

## Materials and methods

### Samples

Whole blood samples of CRC patients from the surgical unit of Aintree University Hospital were collected from May 2022 to November 2022 from patients diagnosed with CRC and about to undergo surgery. White blood cells from 10 ml EDTA-anticoagulated blood were analyzed for immunophenotyping. The immunophenotypic changes were compared between age-matched healthy controls (HCs) and CRCs. The age-matched HCs ranged from 38-89 y (mean ± SD of 59.0 ± 14.1y) and the male: female ratio was 8:9, while the age range of the CRC patients was 34-87y (mean ± SD of 66.8± 13.0 y) and a male: female ratio of 29:20. The study was approved by: the University of Liverpool Committee on Research Ethics (protocol number of UoL001136) for healthy control volunteers; REC reference of 15/NW/0477 (IRAS project ID of 174008) from PINCER (A Platform Study for solId orgaN CancERs) for CRC patients [19]. All participants provided written, informed consent.

### HetaSep separation for peripheral immune cells

Whole blood was mixed with HetaSep solution (Stemcell, USA) at a 5:1 dilution and incubated at 37°C to allow the red cells to sediment about to approximately 50% of the volume. The upper layer was aspirated into a tube containing RPMI 1640 media (Gibco, UK). After mixing, the suspension was centrifuged at 400g for 5 min, the pellet retained and the supernatant discarded. The pellet was gently suspended in 1 mL RPMI medium and then 9 mL of ammonium chloride buffer (13.4 mM KHCO_3_, 155 mM NH_4_Cl, 96.7 µM EDTA) was added and incubated for 4 min to lyse the remaining red cells. After incubation an equal volume of RPMI media was added and immediately centrifuged at 400g for 5 min. The pellet was collected and gently suspended in 1 ml of RPMI media and the cells counted using a cell counter before dilution with RPMI medium to obtain a cell concentration of 0.5 million in 100 μL.

### PBMC preparation

EDTA-anticoagulated blood was layered onto an equal volume of Lymphoprep solution (Stemcell, Germany). The tube was centrifuged at 1000g for 23 min without the break applied. The upper PBMC layer was collected into another tube containing fresh RPMI media and washed at 400g for 5 min. The cell pellet was then suspended in 1 ml RPMI media, and the cells were counted and adjusted with RPMI media to 0.5 million in 100 μL.

### Immunophenotyping

From each sample, 1 million peripheral immune cells were pipetted into the DuraClone immunophenotyping tube (DuraClone IP Phenotyping kit from Beckman Coulter, USA). Each tube contained anti-CD45-KO, anti-CD3-APC Cy7, anti-CD4-APC, anti-CD8-A700, anti-CD19- texa, anti-CD56-PE, anti-CD14-PC7 and anti-CD16-FITC. The added cells were thoroughly mixed by vortexing for 6-8 s and incubated for 15 min at 20°C in the dark. After incubation, the cells were washed with 3ml PBS and then mixed with 500 μL of 0.1% (v/v) formaldehyde before analysis by flow cytometry (BD Fortessa measuring 100,000 events). Compensation with eight single color tubes and QC assessments with standard beads were done before each batch of analysis. FlowJo (v 10.8.1) software was used to analyse the data.

### Surface staining (HLA-DR, CD3, CD14)

PBMC (0.5 million cells/ 100 μL) were washed with 1mL 0.2% (w/v) BSA in PBS and then suspended in 100 μL of 0.2% (w/v) BSA in PBS. Then 1 μL of each fluorescently-tagged antibody (anti-HLA-DR-APC, anti-CD3-PC5.5, anti-CD14-PE) (HLA-DR from Biosciences, USA; CD3, CD14 from BioLegend, USA) was mixed with the cell suspension and incubated at 4°C for 30 min. Excess antibodies were washed out with 1 mL of 0.5M EDTA/PBS and after centrifugation, cells were resuspended in 200 μL of 0.5M EDTA/PBS: 10,000 events were recorded on a flow cytometer (Cytoflex). Compensation was performed for each batch of samples and QC assessment was assessed daily with standard fluorescent beads.

Twenty-eight different types of immune cells subsets were evaluated in this study as shown in Table 1.

**Table 1.** Immune cells subsets of T lymphocytes, NK cells and monocytes

|  |  |  |
| --- | --- | --- |
|  | B lymphocytes | CD45+ CD19+ |
| T lymphocytes subsets | T lymphocytes | CD45+ CD3+ |
|  | Helper T cells | CD45+ CD3+ CD4+ CD8- |
|  | Cytotoxic T cells | CD45+ CD3+ CD4- CD8+ |
|  | Double negative (DN) T cells | CD45+ CD3+ CD4- CD8- |
|  | Double positive (DP) T cells | CD45+ CD3+ CD4+ CD8+ |
|  | Natural killer (NK) T cells | CD45+ CD3+ CD56+ |
|  | CD4+ NK T cells | CD45+ CD3+ CD56+ CD4+ CD8- |
|  | CD8+ NK T cells | CD45+ CD3+ CD56+ CD4- CD8+ |
|  | DN NK T cells | CD45+ CD3+ CD56+ CD4- CD8- |
|  | HLA-DR-positive T cells | CD45+ CD3+ HLA-DR+ |
| NK subsets | NK cells | CD45+ CD3- CD56+/++ |
|  | NK<sup>bright</sup> | CD45+ CD3- CD56++ |
|  |  | CD45+ CD3- CD56++ CD16- |
|  |  | CD45+ CD3- CD56++ CD16+ |
|  | NK<sup>dim</sup> | CD45+ CD3- CD56+ |
|  |  | CD45+ CD3- CD56+ CD16- |
|  |  | CD45+ CD3- CD56+ CD16+ |
|  | CD4+ NK | CD45+ CD3- CD56+/++ CD4+ CD8- |
|  | CD8+ NK | CD45+ CD3- CD56+/++ CD4- CD8+ |
|  | DN NK | CD45+ CD3- CD56+/++ CD4- CD8- |
| Monocytes subsets | Monocytes | CD45+ CD14+/++ |
|  | classical monocytes | CD45+ CD14++ CD16- |
|  | intermediate monocytes | CD45+ CD14++ CD16+ |
|  | non-classical monocytes | CD45+ CD14+ CD16+ |
|  | CD4+ monocytes | CD45+ CD14+/++ CD4+ |
|  | CD56+ monocytes | CD45+ CD14+/++ CD56+ |
|  | HLA-DR<sup>low</sup> monocytes | CD45+ CD14+/++ HLA-DR +(low) |

### Statistical analysis

The normality of data distribution for each immuno-phenotype was tested by the Shapiro-wilk test. The median value, 95% CI, interquartile range (IQR), sensitivity and specificity from ROC were used for presentation of the data. A p value <0.05 is shown as *, and <0.01 as **. Unpaired t-tests and Pearson correlation were applied for normally-distributed data, while Mann-Whitney test and Spearman correlations were applied for skewed data. For comparison of more than two groups, one-way ANOVA for normally-distributed data and Kruskal-Wallis test for skewed data were analysed. Comparisons of cases to control were corrected by controlling the false discovery rate using the two-stages step-up method of Benjamini et al, (2006). GraphPad prism-8 (GraphPad Software Inc., USA) was used for graphical presentations and statistical analyses.

## Results

### **I.** Immunophenotypic changes of CRC patients

Immune cell subsets of T lymphocytes, NK cells and monocytes were compared between age-matched HC and CRC patients. Gating strategies of flow cytometry and immune cell data (mean ± SD) are shown in the supporting information. The IQR of T cells and monocytes was higher in CRC patients, while that of NK cells was lower, and B lymphocytes were significantly lower in CRC patients compared to HC (Fig 1a). Most CRC patients had similar T/B ratios and NK/T ratios compared to HCs, but some patients had either very low or very high T/B ratios and NK/T ratios (Fig 1b, 1c). Although T lymphocytes and monocytes tended to be higher in CRC, these patients had significantly lower L:M ratios (Fig 1d). Most subsets of T cells of CRC patients had higher IQR except DN T cells (Fig 1f), lower numbers of helper T cells but increased numbers of cytotoxic T cells that contributed to lower CD4/CD8 ratios in CRC patients (Fig 1e). The wider IQR of NKT cells in CRC patients (Fig 1f) was related to significant increases of CD4+ NK T (Fig 1h). NK subsets were similar in both groups (Fig 1g), except CD8+ NK cells, which had a wider IQR in CRC patients (Fig 1i). There was a trend for increased numbers of different types of monocytes in CRC patients (increased IQR values) except for classical monocytes which tended to be lower than in HC (Fig 1j, 1k). A significant increase in CD4+ monocytes was observed in CRC patients (Fig 1k).

**Fig 1.**
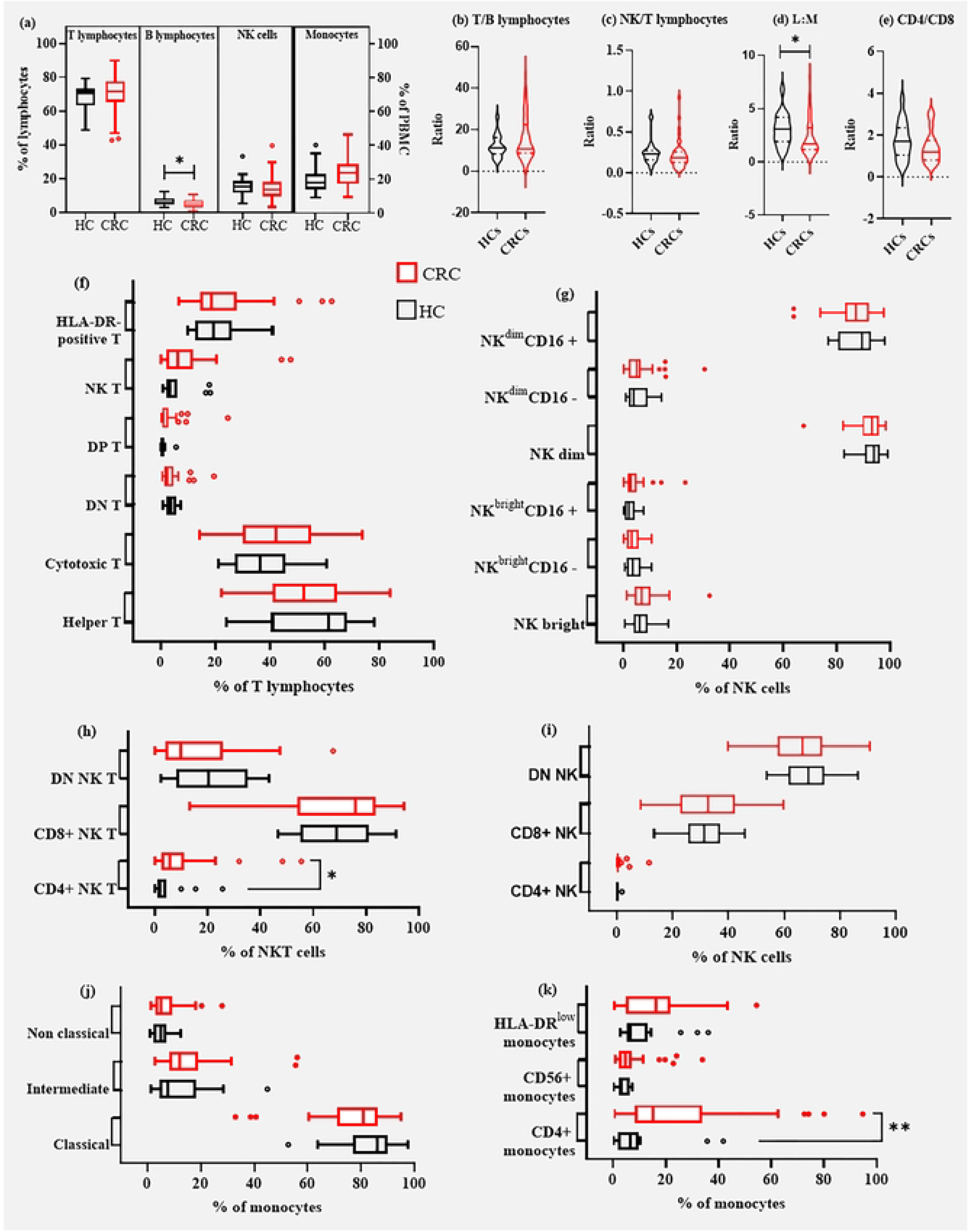
**Comparison of immuno-phenotypes between age-matched HCs and CRCs**. (a) Shows percent of T lymphocytes, B lymphocytes, NK cells and monocytes, with the box representing IQR, line in box represents median, whiskers represent minimum and maximum. (b) Ratio of T lymphocytes to B lymphocytes; (c) ratio of NK cells to T lymphocytes; (d) ratio of lymphocytes to monocytes (L:M); (e) ratio of helper T cells to cytotoxic T cells, all shown as violin plots: solid lines in Violin plots represent median, while dotted lines represent IQR. (f) Helper T, cytotoxic T, DN T, DP T, NK T and HLA-DR positive T lymphocytes. (g) NK^bright^, NK^bright^ CD16-, NK^bright^ CD16+, NK^dim^, NK^dim^ CD16-, NK^dim^ CD16+, (h) CD4+ NK T, CD8+ NK T and DN NK T lymphocytes. (i) CD4+ NK, CD8+ NK and DN NK cells. (j) Classical monocytes, intermediate monocytes and non-classical monocytes. (k) CD4+ monocytes, CD56+ monocytes and HLA- DR^low^ monocytes: all are shown as Tukey graphs and box represents 25% to 75% of samples, lines in box represent median, whiskers represent 3/2 of IQR, dot represents outlier.

### **II.** Immunophenotypic changes in different stages of CRC

Patients’ clinical data showing mean age, male:female ratio and percentage of those >65 y are shown in Table 2. Sixty five percent of CRC patients were above 65 years old and the number of males patients was higher than females. However, there were more females with CRC in the younger population (<50y) and with right-sided tumour cases, and those >65y had increased tumour size, LN involvement and more right-sided tumours.

**Table 2.**
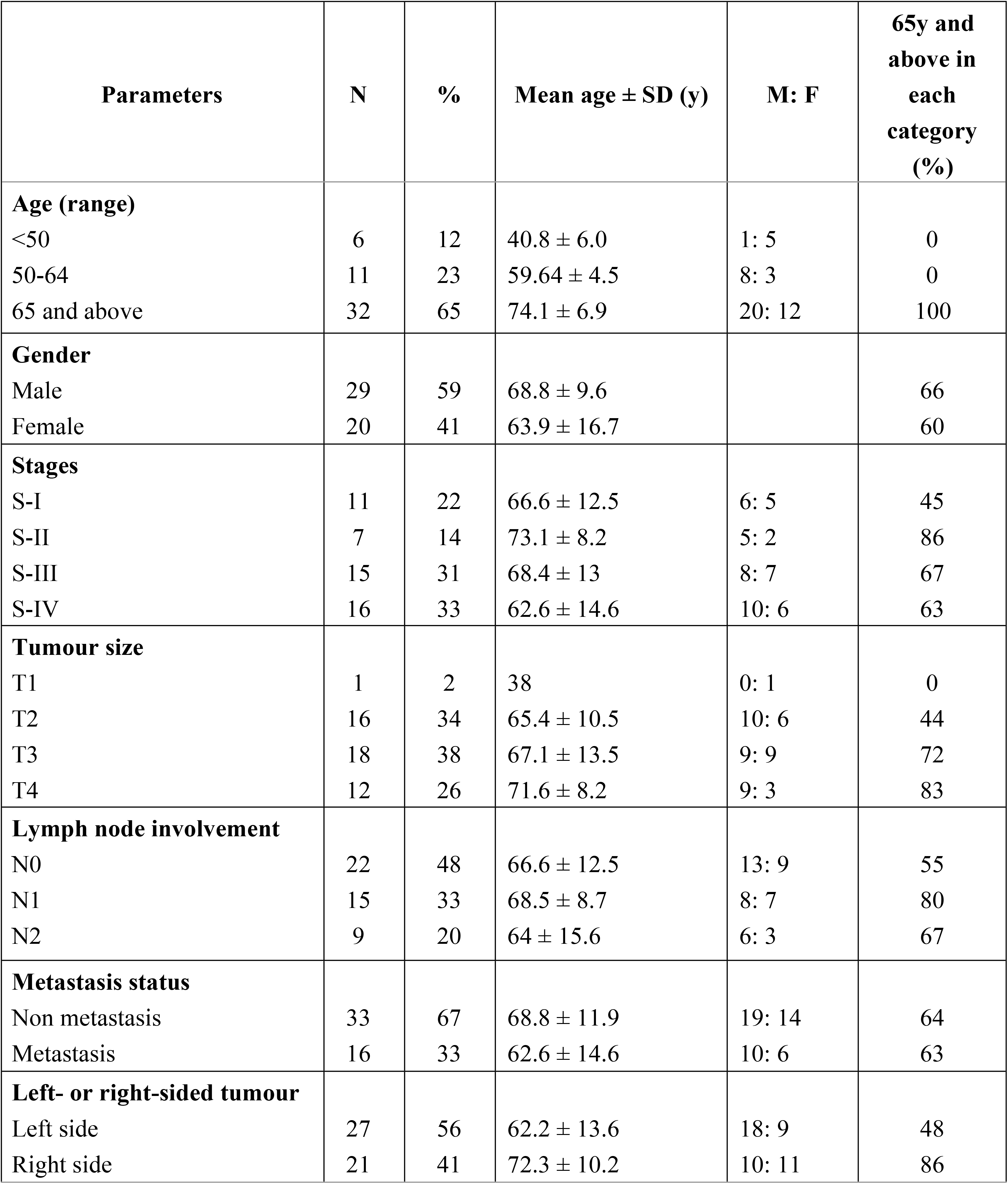
CRC Patients’ demographic data for immunophenotypic study.

| Parameters | N | % | Mean age ± SD (y) | M: F | 65y and above in each category (%) |
| --- | --- | --- | --- | --- | --- |
| **Age (range)** |  |  |  |  |  |
| <50 | 6 | 12 | 40.8 ± 6.0 | 1: 5 | 0 |
| 50-64 | 11 | 23 | 59.64 ± 4.5 | 8: 3 | 0 |
| 65 and above | 32 | 65 | 74.1 ± 6.9 | 20: 12 | 100 |
| **Gender** |  |  |  |  |  |
| Male | 29 | 59 | 68.8 ± 9.6 | | 66 |
| Female | 20 | 41 | 63.9 ± 16.7 | | 60 |
| **Stages** |  |  |  |  |  |
| S-I | 11 | 22 | 66.6 ± 12.5 | 6: 5 | 45 |
| S-II | 7 | 14 | 73.1 ± 8.2 | 5: 2 | 86 |
| S-III | 15 | 31 | 68.4 ± 13 | 8: 7 | 67 |
| S-IV | 16 | 33 | 62.6 ± 14.6 | 10: 6 | 63 |
| **Tumour size** |  |  |  |  |  |
| T1 | 1 | 2 | 38 | 0: 1 | 0 |
| T2 | 16 | 34 | 65.4 ± 10.5 | 10: 6 | 44 |
| T3 | 18 | 38 | 67.1 ± 13.5 | 9: 9 | 72 |
| T4 | 12 | 26 | 71.6 ± 8.2 | 9: 3 | 83 |
| **Lymph node involvement** |  |  |  |  |  |
| N0 | 22 | 48 | 66.6 ± 12.5 | 13: 9 | 55 |
| N1 | 15 | 33 | 68.5 ± 8.7 | 8: 7 | 80 |
| N2 | 9 | 20 | 64 ± 15.6 | 6: 3 | 67 |
| **Metastasis status** |  |  |  |  |  |
| Non metastasis | 33 | 67 | 68.8 ± 11.9 | 19: 14 | 64 |
| Metastasis | 16 | 33 | 62.6 ± 14.6 | 10: 6 | 63 |
| **Left- or right-sided tumour** |  |  |  |  |  |
| Left side | 27 | 56 | 62.2 ± 13.6 | 18: 9 | 48 |
| Right side | 21 | 41 | 72.3 ± 10.2 | 10: 11 | 86 |

Changes in numbers of immune cells in patients at clinical stages of CRC (assessed by the American Joint Committee on Cancer TNM system) were determined and a decreased L:M was observed throughout the stages of CRC that was statistically-significant in stage 2 and stage 3. There was a trend for a decrease in CD4/CD8 in late stages of CRC (Fig 2a). Levels of DP T cells were significantly increased in stage 2 and stage 3 (Fig 2b). Increased levels of NK T cells related to the increased levels of CD4+ NK T cells and the CD8+ NK T cells in all stages of CRC (Fig 2c). No obvious changes were observed in NK cells subsets (Fig 2d, 2e, 2f). There were changes of monocytes subsets at different stages of CRC with a trend for decreases of classical monocytes, increases of intermediate monocytes and statistically-significant increases of CD4+ monocytes (Fig 2g).

**Fig 2.**
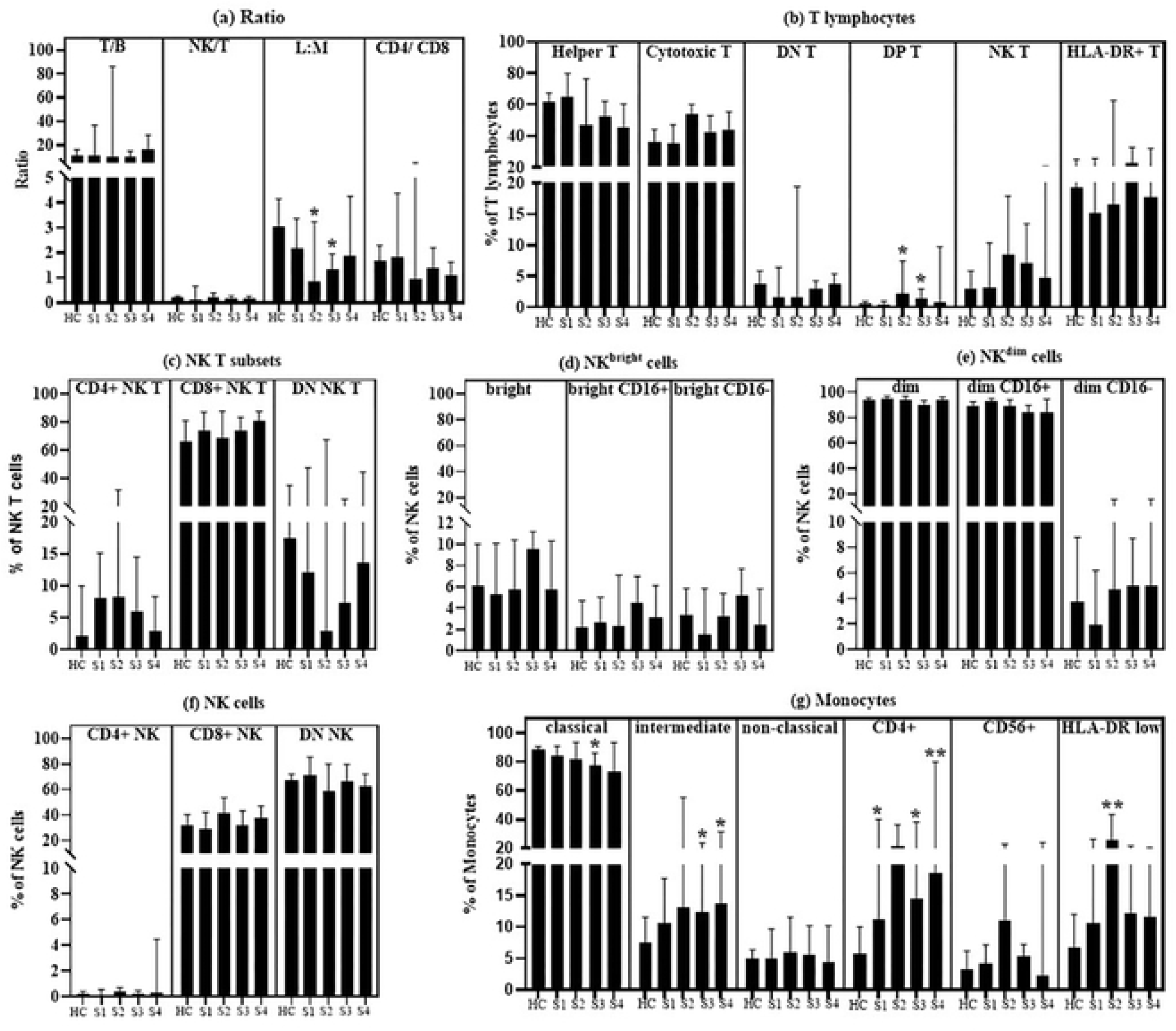
Changes of immuno-phenotypes during the clinical stages of CRC. (a) Ratio of T lymphocytes to B lymphocytes, NK cells to T lymphocytes, lymphocytes to monocytes ratio (L:M) and helper T cells to cytotoxic T cells. (b) Helper T, cytotoxic T, DN T and DP T, NK T and HLA-DR positive T. (c) Different types of NK T cells (CD4+ NK T, CD8+ NK T and DN NK T). (d) NK^bright^ cells (total NK^bright^, NK^bright^ CD16+ and NK^bright^ CD16-). (e) NK^dim^ cells (total NK^dim^, NK^dim^ CD16+ and NK^dim^ CD16-). (f) Different types of NK cells (CD4+ NK, CD8+ NK and DN NK). **(g)** Different types of monocytes (classical, intermediate and non-classical), CD4+ monocytes, CD56+ monocytes and HLA-DR^low^ monocytes. Bars represent median values, whiskers represent 95% CI.

### **III.** Immunophenotypes relating to tumour sizes and LN involvement

Immune cells function in a co-ordinated manner in their role against tumours or microbial pathogens [20] and tumours can alter the behaviour of the immune cells to favor their survival and progression [21]. We therefore measured immunophenotypes in CRC patients and determined if they were correlated with tumour size (T1-T4) or LN involvement. Most immunophenotypes showed no correlation with tumour sizes or LN involvement. However, subsets of NK cells showed either positive correlation or negative correlation with tumour sizes. For example, NK^dim^ cells, especially NK^dim^ CD16+ cells, which are the highest population of NK subsets, were significantly decreased while NK^bright^ cells were increased in patients with large tumours (Table 3).

**Table 3.** Immunophenotypes relating to tumour sizes and LN involvement

| **Immunophenotypes**<br>**negatively correlated**<br>**(decreased) with tumour**<br>**size** | **r** | **p** | **Immunophenotypes**<br>**positively correlated**<br>**(increased) with tumour**<br>**size** | **r** | **p** |
| --- | --- | --- | --- | --- | --- |
| NK<sup>dim</sup> CD16<sup>+</sup> | - 0.5 | 0.002 | NK<sup>dim</sup> CD16<sup>-</sup> | 0.4 | 0.02 |
| NK<sup>dim</sup> | - 0.3 | 0.04 | NK<sup>bright</sup> | 0.3 | 0.04 |
| DN NK | - 0.3 | 0.07 | DP T | 0.3 | 0.07 |
|  |  |  | NK<sup>bright</sup> CD16<sup>+</sup> | 0.3 | 0.1 |
|  |  |  | CD8<sup>+</sup> NK | 0.3 | 0.1 |
| **Immunophenotypes**<br>**negatively correlated**<br>**(decreased) with LN**<br>**involvement** | **r** | **p** | **Immunophenotypes**<br>**positively correlated**<br>**(increased) with LN**<br>**involvement** | **r** | **p** |
| NK<sup>dim</sup> CD16<sup>+</sup> | - 0.3 | 0.06 | NK<sup>bright</sup> CD16<sup>-</sup> | 0.3 | 0.08 |

### **IV.** CD4+ monocytes in CRC

CD4+ monocytes were significantly higher in CRC patients compared to HC (Fig 1k). In addition, CD4+ monocytes were positively correlated with the numbers of intermediate monocytes, NK^dim^ cells, CD4+ NK cells and DP T cells, but negatively correlated with numbers of classical monocytes, NK^bright^ and DN T (Fig 3a). These changes are likely to be related to numbers of intermediate monocytes, because intermediate monocytes had an increased level of CD4 staining intensity among the three subsets of monocytes, and CRC patients had significantly-increased CD4 staining intensity of intermediate monocytes compared to HC (Fig 3b). We also analysed paired samples of similar age and same sex, e.g., 59y female HC and 59y female CRC. There were 15 pairs of participants where we could perform this age- and sex-match (M: F ratio of 7:8) and CD4+monocytes were significantly-increased in CRCs (p=0.003) after this paired data analysis (Fig 3c). CRC and HC data for CD4+ monocytes were further analysed by receiver operating characteristic (ROC) analysis to determine if their numbers have potential as a biomarker to indicate CRC. The area under the curve was 76% (p=0.002) and the standard error was 0.07: there was 60% sensitivity and 88% specificity with a likelihood ratio of 4.8, cut-off value of 11.6% (Fig 3d, 3e).

**Fig 3.**
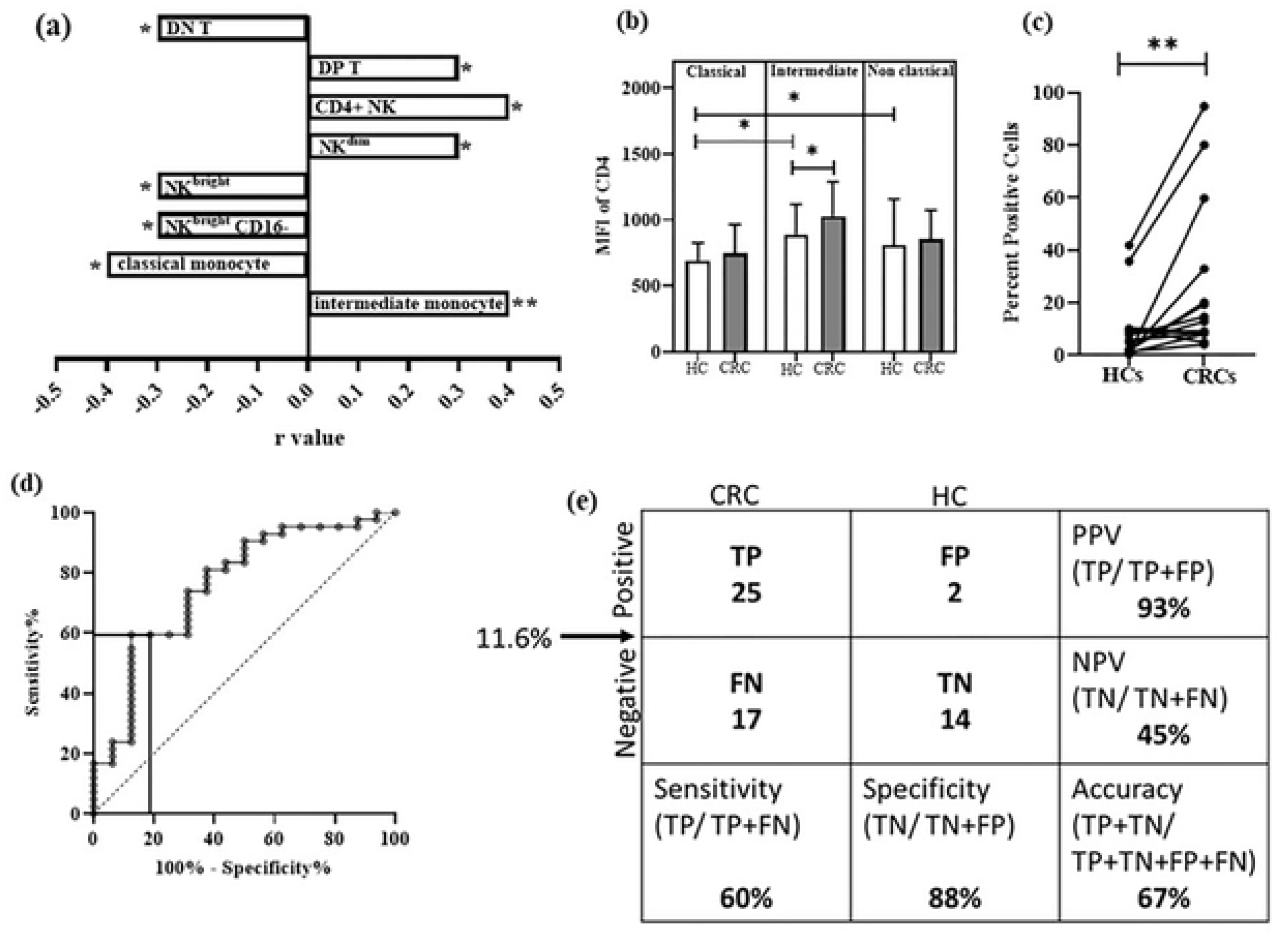
Analysis of CD4+ monocytes. (a) Shows positive and negative correlations with other immunophenotypes. (b) CD4 staining intensity expressed by classical, intermediate and non-classical monocytes in age-matched HCs and CRC. (c) Comparison of CD4+ monocytes in paired samples of HCs and CRCs. (d) ROC analysis of CD4+ monocytes between age-matched HCs and CRCs, the cut-off value of 11.6% at the vertical solid line and horizontal solid line showing 60% sensitivity and 88% specificity. (e) 2x2 table of CD4+ monocytes showing sensitivity, specificity, positive predictive value (PPV), negative predictive value (NPV) and accuracy. TP = true positive, FP = false positive, FN = false negative, TN = true negative.

## Discussion

One of the hallmarks of cancer is evasion of immune system. Immune cells are a heterogenous and dynamic population of cells and are involved in the surveillance of tumour cells, immune responses to cancers, eradication of cancers and progression of cancer [22]. However, similar changes in immune cells may occur during ageing and because incidence of CRC increases with age, it is important to distinguish immune changes that are specific to CRC rather than those that develop as a function of age. In cancers, some changes in immune cells can aid recognition and destruction of the tumours, while other changes can impair immune surveillance and allow the tumour cells to avoid recognition and destruction. In this report, we show changes in specific immune cell subtypes in CRC patients that are not seen in age-matched HC. While we have measured changes in numbers of individual immune cell subtypes, effective immune responses requires co-ordinated interactions for both positive and negative regulation of immune function. Thus, changes in numbers or function of one cell type can have profound effects of the regulation of activity of other immune cells. For example, coordination between T cells and B cells is important for successful T cell-dependent antibody responses [23]. In addition, communications between NK cells and T cells are important for the initial regulation of T cells [24]. One striking finding of our study was the lower L:M ratio that was seen throughout the stages of CRC indicating there was a myeloid skew in CRC patients even in the early stages of this disease.

### T lymphocytes

Most CRC patients had decreased numbers of helper T cells and increased cytotoxic T cells. This could negatively impact on the ability of helper T cells to regulate cytotoxic T cells to mediate their effector functions through CD27 co-stimulation. With lower numbers of helper T cells, cytotoxic T cells may become exhausted by the upregulation of PD-1 and co-inhibitory receptors [25]. Therefore, decreased helper T cells can negatively affect the cytotoxic T cells and “helpless” cytotoxic T cells can have impaired anti-tumour activity. Most CRC patients had lower CD4+ T/ CD8+ T cells and this ratio has been used as a prognostic biomarker with a lower ratio indicates tumour progression [26]. Moreover, decreased trends of helper T cells and increased cytotoxic T cells were observed starting from SII of CRC. Therefore, there may be exhausted cytotoxic T cells in CRC even at the early stages of the disease.

### NK T cells

NK T cells can cross-talk with both innate and adaptive immune cells, but exist in different subsets that can have either pro-tumour or anti-tumour properties [27, 28]. The majority of CRC patients had higher numbers of total NK T cells, particularly in the subsets of CD4+ NK T. The expression of co-receptors on NK T cells relates to immunosuppression: these T cells are involved in CD8+ Treg generation while CD8+ NK T cells are involved in the immune response by controlling antigen-bearing DCs [29, 30]. On the other hand, DN NK T cells have Th1-like function, and they can kill tumour cells [28]. They were present in lower numbers in most CRC patients and so an increased number of NK T cells in CRC indicates a phenotype of immunosuppression rather than anti-tumour activity.

### DN T, DP T cells

Although DN T cells (CD3+ CD4- CD8-) and DP T (CD3+ CD4- CD8-) are present in low numbers in the bloodstream, they can have distinct and complex functions. For example: DN T cells may have immunoregulatory roles, e.g. to prevent graft-versus-host-disease [31]; DP T cells are activated cytotoxic cells [32] and can have distinct complex functions because they are positive for both CD4 and CD8 receptors. These latter cells have been shown to have anti-tumour as well as pro-tumour activities in tumour immunity [33]. Most CRC patients had lower levels of DN T cells, but had higher numbers of DP T cells, especially in SII and SIII, indicating complex immune cell functions involved in the progress of CRC.

### NK cells

NK cells are considered cytotoxic cells of innate immunity and are important not only for the killing of tumour cells but also for tumour remission [34]. It has been shown that increased numbers of NK cells relate to higher survival rates in colorectal cancer patients [35]. In this study, the total NK cells count tended to decrease in CRC, but phenotypic subsets were similar to HCs. However, NK cells showed a correlation with tumour size: NK^bright^ were increased and NK^dim^ decreased especially NK^dim^ CD16+ cells as tumour size increased. NK^bright^ cells are considered immature cells having increased production of cytokines such as IFN-γ, TNF-α, GM-CSF, IL10 and IL13 and have less cytotoxic activity than NK^dim^ cells [36]. The numbers of NK^bright^ cells were increased in larger tumours implying that while they have decreased cytotoxic function, they may be capable of regulating the immune response via their cytokine production. However, the major population of NK cells, NK^dim^ CD16+ decreased which may also indicate decreased ability to target tumour cells, as NK^dim^ CD16+ are important for the cytotoxicity of NK cells [36]. Therefore, these immunophenotypic changes of NK cells in CRC could lead to inefficient tumour cell clearance in CRC.

### Monocytes

Most patients with CRC had lower numbers of classical monocytes, higher numbers of intermediate monocytes, with increased numbers of non-classical monocytes. This result was similar to that of Schauer et al (2012) who showed that intermediate monocytes were increased in CRC patients, especially in localized tumour cases. They predicted that the intermediate monocytes could be used as diagnostic markers, with 69% sensitivity and 81% specificity in CRC [37]. During progression of the disease (SI-IV), most CRC patients had increased intermediate monocytes throughout all stages, but non-classical monocytes increased in S-II and decreased in later stages. The effector function of immune cells, as well as their ability to phagocytose and clear debris, should be in balance for a successful immune response. Defects in clearance functions can lead to tumour progression especially through the inflammation process [38]. As non-classical monocytes have a role in patrolling and scavenging debris, their decreased levels could relate to impaired clearance of debris in response to the tumour.

HLA-DR positivity of monocytes changed in CRC, with the majority of CRC patients showing increased numbers HLA-DR^low^ monocytes. HLA-DR expression is important for antigen presentation by monocytes and so impaired expression of HLA-DR may indicate a defect in this function of monocytes. In contrast, HLA-DR^low^ positivity represents immunosuppression [39]. CD56+ monocytes increased in most CRC patients and CD4+ monocytes increased significantly. CD56 expression relates to T cell proliferation [40], while CD4 expression relates to differentiation of monocytes [41]. Therefore, monocytes of CRC were functionally diverse: they could be functionally active as well as immunosuppressive in the immune response of CRC patients. During disease progression, CD4+ monocytes and CD56+ monocytes showed increased trends in S-II and S-III. However, CD56+ monocytes showed a decreased trend in S-IV. Moreover, the immunosuppressive phenotypes HLA-DR^low^ monocytes significantly increased in S-II. Therefore, the functional properties of monocytes were changing throughout the stages of the disease.

An increased positivity of CD4 was observed in monocytes as well as in NK cells (there was a positive correlation between cell numbers of CD4+ monocytes and CD4+ NK cells). The increased levels of CD4+ monocytes likely related to increased levels of mature cells and more complex cells (NK^dim^ and DP T) plus decreased levels of immature and immunoregulatory cells (NK^bright^ and DN T). It may also be connected with numbers of intermediate monocytes: levels of CD4+ monocytes and intermediate monocytes increased, but decreased levels of classical monocytes were observed in CRC patients. In addition, CD4 staining intensity was significantly increased in intermediate monocytes compared to classical monocytes in CRC patients compared to age-matched HCs. This increase was observed when all pooled CRC and HC samples were compared, and this increase was even more significant in age- and sex-matched CRC/HC pairs. When CD4+ monocytes were evaluated as a potential diagnostic marker in CRC, the analysis showed a 60% sensitivity and 88% specificity with the cut-off value of 11.6%. Based on data in the literature and the findings of the study, the schematic diagram in Fig 4 summarises the major changes in monocytes during ageing and CRC progression. We propose that increased CD4+ monocytes could relate to and increased workload of monocytes due to ageing and CRC.

**Fig 4:**
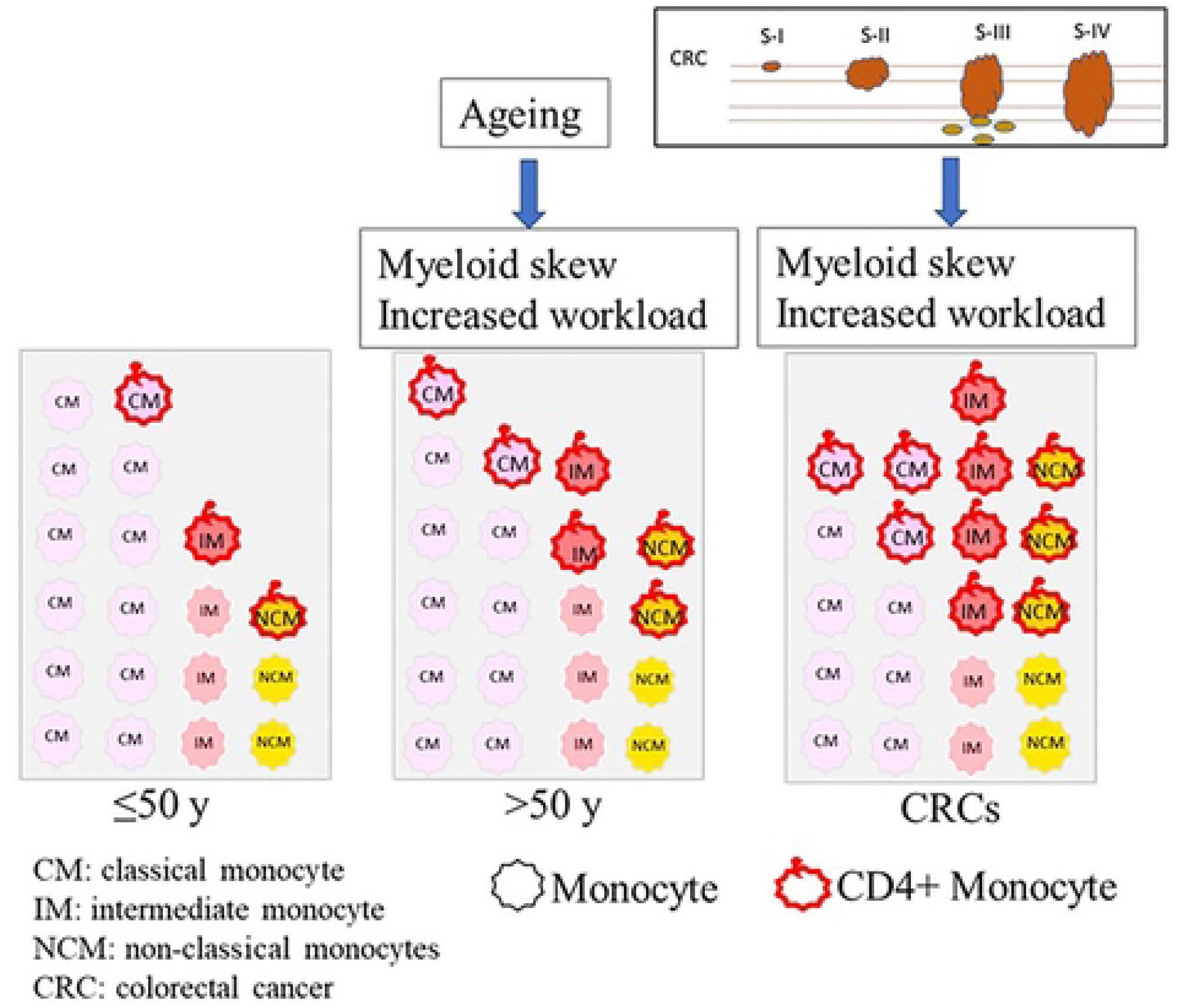
Possible explanation of increased CD4+ monocytes in ageing and CRC

In conclusion, we show complex changes in the immune phenotypes in CRC that are distinct from those that occur during ageing, and many of these changes are in immune cells subtypes that promote an immuno-suppressive phenotype. These changes may allow the tumour cells to evade immune surveillance and targeted destruction. The significantly increased level of CD4+ monocytes could be a potential diagnostic marker for CRC patients.

## List of abbreviations

CRC: colorectal cancer
DN: Double negative
DP: Double positive
HC: Healthy control
IQR: Interquartile range
L:M: lymphocytes to monocytes ratio NK Natural killer
NPV: negative predictive value
PBMC: Peripheral blood mononuclear cells PPV positive predictive value
ROC: receiver operating characteristic

## Acknowledgements

We express our gratitude to Asso. Prof. Jaranit Kaewkungwal from the Faculty of Tropical Medicine, Mahidol University for assistance with the statistical analyses.

## Supporting information

S1 Fig. Gating strategies of immunophenotypes

S1 Table. Comparison of immunophenotypes (percent positive cells) in age-matched HCs and CRC patients

